# Tumoroid-on-a-Plate (ToP): Physiologically Relevant Cancer Model Generation and Therapeutic Screening

**DOI:** 10.1101/2024.04.15.589651

**Authors:** Amir Seyfoori, Kaiwen Liu, Hector Caruncho, Patrick Walter, Mohsen Akbari

## Abstract

Employing three-dimensional (3D) *in vitro* models, including tumor organoids and spheroids, stands pivotal in enhancing cancer therapy. These models bridge the gap between 2D cell cultures and complex in vivo environments, effectively mimicking the intricate cellular interplay and microenvironmental factors found in solid tumors. Consequently, they offer versatile tools for comprehensive studies into cancer progression, drug responses, and tailored therapies. In this study, we present a novel open-surface microfluidic-integrated platform called the Tumoroid-on-a-Plate (ToP) device, designed for generating intricate predictive 3D solid tumor models. By incorporating a tumor mass, stromal cells, and extracellular matrix components, we successfully replicate the complexity of glioblastoma (GBM) and pancreatic adenocarcinoma (PDAC) within our system. Using our advanced ToP model, we were able to successfully screen the effect of various GBM extracellular matrix compositions, such as Collagen and Reelin, on the invasiveness of the GBM cells with the ToP model. The ToP in vitro model also allowed for the screening of chemotherapeutic drugs such as temozolomide and iron-chelators in a single and binary treatment setting on the complex ECM-embedded tumoroids. This helped to investigate the toxic effect of different therapeutics on the viability and apoptosis of our in vitro GBM and PDAC cancer models. Additionally, by co-culturing human-derived fibroblast cells with PDAC tumoroids, the pro-invasive impact of the stromal component of the tumor microenvironment on growth behaviour and drug response of the tumoroids was revealed. This study underscores the transformative role of predictive 3D models in deciphering cancer intricacies and highlights the promise of ToP in advancing therapeutic understanding.

## 1- Introduction

Three-dimensional (3D) cell culture modelling is an indispensable tool for *in vitro* efficacy testing in the drug development process. New fabrication methods for producing uniform and large quantities of multicellular spheroids (MCS) or organoids present a promising solution to costly animal testing and predictive shortcomings in monolayer or “2-dimensional” *in vitro* cultures^1–3^. Compared to traditional monolayer cultures, 3D culture models better mimic the cellular microenvironment, intercellular crosstalk, susceptibility to drugs, and structural morphology of *in vivo* tissue, which is highly desirable when transitioning to animal testing and clinical trials ^1,2^. In addition, these models can gain additional complexity and physiological relevance through the introduction of different bioengineered extracellular matrices (ECM). 3D in-vitro models that involve ECM provide bio-mimicked microenvironments that replicate various signalling transduction processes occurring in cell-cell and cell-ECM interactions. This will structurally and biochemically support the tumour organoid growth and maturity ^2,4,5^. Multiple studies have demonstrated that creating a 3D model of the intricate tumor microenvironment can enhance tumor spheroids’ ability to simulate a conducive environment for testing the effectiveness of therapeutic agents. This process also emphasizes the crucial cell-matrix interactions that result in invasion and malignancy ^4,6,7^. As a result, 3D cell culture spheroid/organoid models may provide a high-throughput drug screening solution for anti-tumor drug testing and be more predictive in clinical translations^6,7^.

Simple 3D tumor models using spheroids can be generated from a broad range of adherent cell types including patient-derived tumor and cancer cell lines. Spheroids are formed via self-organization and intercellular linking of cancer cell lines through cadherin and integrin binding and, as such, reconstitute physiological structures and signaling pathways that closely resemble the corresponding tissue *in vivo* ^8,9^. Patient-derived organoids are very similar to cancer spheroids but encompass a collection of organ- or tissue-specific cell types produced from patient progenitor and stem cells, yielding structures and functions exceedingly similar to tissues and organs ^10,11^. Patient-derived organoid and spheroid models have contributed tremendously to cancer research and spearheaded advancements in personalized medicine due to their drug discovery and precision medicine applications ^11,12^. Patient-derived organoid models are particularly advantageous as a drug testing platform because of their phenotype-specific copying of patient organs and ability to reproduce genetic heterogeneities of in vivo tumors ^10,13^. Furthermore, modelling these patient-derived cells with their unique genetic signatures creates more realistic tumor responses to particular metabolic demands of chemotherapy or environmental manipulations such as hypoxia. Adaptations of these models for high throughput screening have opened the possibility to test organoid and spheroid responses to numerous combinations and dosages simultaneously, increasing treatment accuracy in the clinic and highlighting their potential for personalized medicine ^9,13^.

There are several methods to produce multicellular spheroids based on the non-cell adherent substrates, including soft lithography, 3D printing (stereolithography), injection molding, and thermoforming microwells^15,16^. Among these fabrication methods, hydrogel microwells generated by 3D printing are particularly well suited for high throughput drug testing applications due to their high precision, versatility, and low cost ^17^. Microfluidic devices that rely on soft lithography techniques are often time-consuming to construct as the process involves several labor-intensive steps and require specialized equipment. Hydrogel microwells fabrication using molds from 3D resin printers addresses these issues by being fast, accurate, and highly scalable ^18^. Multiple replicas of the same mold can be printed, allowing for large-scale production of microwells and flexibility when prototyping different designs. Additionally, hydrogel microwells have been proven to maintain high cell viability when forming spheroids due to the bio-mimetic properties of hydrogels and offer great control over the size of spheroids ^19,20^. Several studies have employed a variety of gels to create microwell platforms such as agarose^1,18^ chitosan^19^, and gelatin methacryloyl (GelMA) ^21,22^. Among these, agarose emerged as an attractive option for a microwell hydrogel due to its biocompatibility, non-adherent nature, and structural characteristics.

The current hydrogel-based microwell arrays have limitations in creating a comprehensive environment that can fully replicate all the different compartments of the tumor microenvironment, including the ECM and stromal compartments. Therefore, there is an unmet need to develop a new platform that can overcome these limitations and provide holistic modeling of the tumor microenvironment.

To tackle these concerns, we have developed a platform called tumoroid on-a-plate (ToP) that enables the creation of 3D tumor spheroids/organoids within the hydrogel inserts inside the standard well plates. This allows for the study of important cellular parameters and drug responses, because of the replication of complex cellular microenvironments such as realistic cellular compositions and ECM environments. ToP is scalable and high-throughput and features tailored matrices for different disease models. It also allows for efficient assay evaluation on the same plate. The ToP platform builds upon our previous self-filling microwell arrays (SFMAs) technology, which produces many uniform tumor spheroids using an inclined surface and micro-channels that distribute cells evenly to an array of microwells with the help of gravity^18^. Our current work uses a prefabricated agarose scaffold with a unique design and geometry for rapid spheroid formation and enables efficient deposition of different ECMs for studies that include angiogenesis, invasiveness, morphogenesis, and malignancy. Currently, operational procedures for conventional spheroid–embedded ECM models are labor-intensive and time-consuming because the relocation of the spheroid and replacement of the medium is required ^5^. Oftentimes, multiple spheroids must be physically manipulated, and several studies have reported well-to-well variation and spheroid damage during the relocation process ^15,23,24^. On this basis, our ToP platform is designed to offer a dynamic and versatile model that can perform three tasks in one insert. This includes: 1) creating uniform spheroid construction, 2) enabling facile co-culture by applying ECM and other tumor microenvironment (TME) components within the well, and 3) conducting drug testing and downstream analysis without the need for spheroid transfer. We utilized the current platform to create glioblastoma tumor models and investigate the impact of Reelin protein on the GBM microenvironment. Specifically, we studied how this protein could affect tumour invasion behavior when combined with single or multiple therapeutic agents such as Temozolomide and deferoxamine. Additionally, the current ToP platform was used to model the role of stromal cells in the microenvironment of the pancreatic tumor. We showed that the current platform could incorporate stromal fibroblast cells in both direct and indirect co-culture methods in separated compartmentalized microchannels without needing to transfer and manipulate the generated tumoroids in the plate.

## 2- Experimental

### 2-1- Materials

Agarose was purchased from Bio Basic Inc. (Markham ON, Canada). Dulbecco’s Modified Eagle Medium (DMEM), fetal bovine serum (FBS), Dulbecco’s phosphate-buffered saline (DPBS), and penicillin/streptomycin (pen/strep) were purchased from Gibco (Grand Island, NY, USA). 6-Well plate was obtained from Corning (Corning, NY, USA). High-resolution photo-curable resin was purchased from AnyCubic (Shenzhen, Guangdong, China). Temozolomide, gemcitabin and live/dead staining viability kit, were purchased from Millipore Sigma (Oakville, ON, Canada). The 10× phosphate buffered saline (PBS), 0.5 N NaOH and bovine collagen type 1 (10 mg/mL) were purchased from Advanced BioMatrix Inc. (San Diego, CA, USA). Recombinant Human Reelin Protein was purchased from the R&D systems Inc. (Minneapolis, USA).

### 2-2- 3D printing of mold and hydrogel microwell fabrication

The high throughput model-making platforms were fabricated by casting molten agarose onto a digital light processing (DLP) 3D printed mold. The mold was designed using SOLIDWORKS (SolidWorks Corp., USA) and was exported to an STL format for printing. The STL file was sliced at a 15 μm thickness and printed using an AnyCubic Photon S 3D printer with a high precision ultraviolet (UV) sensitive polymer resin. Each platform contained 24 microwells separated into 4 quadrants that were connected to cell loading areas through guiding microchannels. The U-shaped microwells had a diameter of 700 μm. Once the mold was printed, 2% wt/vol agarose in PBS solution was prepared by heating it up to its boiling temperature and then allowed to cool to 60 °C before it was cast onto the mold. After 8 minutes, the now gelled agarose microwell insert was removed from its mold and transferred to a sterile 6-well plate. The microwell insert and the 6-well plate were sterilized under UV light with a maximum wavelength of 365 nm for 3 hours.

### 2-3- Tumoroid formation and ECM embedding

Human-derived glioblastoma cells, U251, were cultured in DMEM supplemented with 10% FBS, 100 IU ml−1 penicillin and 100 mg ml^−1^ streptomycin. U251 multicellular tumor spheroids formed by seeding 2×10^5^ cells onto the loading zone for each quadrant. The cells were incubated at 37C, and ten minutes was allowed to settle, slide down the guiding channels, and fill the microwells. After, the excess cells were aspirated from the channels and washed with fresh media twice. The U251 spheroids were monitored daily to assess spheroid formation and their growth over three days.

The 3-in-1 culture plate has an open-surface design which allows easy ECM hydrogel embedding without the need to transfer the tumoroids. The glioblastoma ECM microenvironment was mimicked using bovine fibril collagen with a working concentration of 4 mg/mL. The hydrogel solution was prepared by adding 10X PBS and 0.5 N NaOH to the stock solution of collagen with a ratio of 1:1:8. The hydrogel solution was then diluted with culture media until the concentration reached 4 mg/mL. To assess the effects of reelin recombinant protein on glioblastoma survival and invasion, stock Reelin solution was mixed with the dilution culture media to a concentration of 10 nM to produce the reelin condition. The final collagen solution was gently dispensed onto the ECM loading zone and then incubated at 37°C for 1 hour to complete cross-linking of the hydrogel.

### 2-4- Tumoroid drug testing

The Glioblastoma tumoroids were treated with different combinations and concentrations of temozolomide (TMZ) and deferoxamine (DFO). Four-day old tumoroids embedded in collagen-Reelin ECM were treated first with DFO at 10 µM and 100 µM concentrations and exposed for three days. The tumoroids were treated without the need for relocation. The primary culture media was aspirated before simply applying the treatment-containing media afterwards to the media reservoirs. For the combinational drug treatment study, after three days of DFO treatment, TMZ with different concentrations of 50,100,250 and 500 μM in culture media was added to the tumoroids. The control samples for comparison were tumoroid growth without any treatment, tumoroids treated with 10 µM and 100 µM only and tumoroids treated with 500 μM TMZ only. Tumoroids of the pancreas were treated with Gemcitabine (GEM) and DFO in monoculture and co-culture settings. The tumoroids were embedded in collagen ECM and were treated with varying concentrations of GEM (1-1000 µM) and DFO (10-100 µM) for three days. The treatment duration was the same as that for GBM tumoroids.

### 2-5- Metabolic activity assay of the tumoroids after drug testing

Presto Blue viability assay was used to quantify drug toxicity of different doses and combinations of drugs. This assay measures the metabolic activity of live tumor cells and has fluorescence excitation and emission wavelengths of 560 nm and 590nm, respectively. To prepare the Preto Blue assay, the Presto Blue reagent was added to fresh culture media to create a 10% vol/vol solution and was added directly to the media reservoir of the insert and over the tumoroids. The entire setup was incubated for 4 hours, after which, samples were placed into a plate reader. The average fluorescent intensity of the spheroid’s samples was subtracted from the intensity of a blank cell-free sample.

### 2-6- Tumoroid cell staining on-a-plate

To assess the viability of the tumor cells withing the spheroids, the Live/Dead (L/D) assay was conducted using 1 μM calcein AM and 4 μM ethidium homodimer-1 (Life Technologies kit) for 1 hour at 37 °C. The assay was applied to the entire ECM-embedded glioblastoma tumoroid model and imaged on the culture plate without the need to remove the tumoroid. The effects of drug dosage and combinational drug treatment were assessed by fluorescent imaging and the invasion length of invaded cells with the ECM was quantified using ImageJ.

To perform fibronectin and E-cadherin immunostaining of the tumoroids within the plate, they were first fixed with 10% neutral buffered formalin (Thermo Fisher Scientific, Waltham, USA) for 45 minutes at room temperature. After that, they were washed three times with PBS. The fibers were then permeabilized with 0.3% Triton-X100 in PBS for 10 minutes, followed by incubation in a blocking buffer of 3% BSA and 0.3% Triton-X100 in PBS for 30 minutes. Then, the blocking buffer was removed, and DPBS was added to the reservoirs of the insert and incubated overnight at 4 °C in the dark. To prepare the antibody solution, the fluorescent tag-conjugated Fibronectin (PE) and E-cadherin (FITC) antibodies were diluted in 1% BSA and 0.3% Triton X-100 in DPBS solution to a specific concentration. The solution was added to the secondary reservoir of the insert and the plate was incubated for 2-4 hours at 4 °C in the dark. Before immunofluorescence imaging, the samples were washed three times with DPBS.

### 2-7- Real-time PCR analysis

To determine whether 3D cultured tumoroids responds differently to the treatment of different concentrations of iron chelator and TMZ (IC50) and possible apoptosis induction, real-time PCR performed for pro/anti-apoptotic genes bcl-2 and Bax. With this regard, mRNA extraction and DNase digestion were performed following the instruction of Qiagen’s RNeasy Mini Kit and RNase-Free DNase kit, respectively. Extracted-RNA purity was tested using Nandrop. First-strand cDNA was synthesized following Invitrogen’s SuperScript III First-Strand Synthesis System, and the purity was tested by Nandrop. The cDNA was used as a template for real-time PCR. Primers designed to specifically amplify cDNA from bax and bcl-2 mRNAs (GAPDH used as housekeeping/internal control). QuantiNov SYBR Green PCR Kit was then used to prepare the reactions for quantitative measurement of bcl-2 and Bax genes.

### 2-8- Statistical analysis

In each experiment, we used a minimum of three repetitions and presented the data as mean ± standard deviation (SD). To compare the means, we performed a one-way ANOVA with Tukey’s multiple comparisons using GraphPad Prism 7.0. We considered the following values for statistical significance: *p < 0.05, **p < 0.01, ***p < 0.001, and ****p < 0.0001.

## 3- Result and Discussion

### 3-1 Fabrication and Characterization of tumoroid-on-a-plate (ToP) Model

To mimic the complexity of the tumor microenvironment, including growth in 3D multicelluer spheroid, tumor-associated stroma, and extracellular matrix, we developed a microfluidic-integrated culture plate where the molecular crosstalk between different compartments of the TME impacts the invasiveness of the tumor due to modelling in physiologically relevant environments. The developed platform can recapitulate the overall steps of tumor initiation, progression and migration in a universal platform without the need for extra manipulation of the formed tumoroid and well-to-well transferring. We recreated this configuration using the agarose-based insert in-a-well with open surface inclined microchannels connected to the self-filling microwells, Fig. 1A&B. The hydrogel insert comprises four quadrants, enabling four different parallel experiments in a single well. In each quadrant, there are six microwells, which are addressable with secondary microchannels to a reservoir, enabling exposing the tumoroids to therapeutic compounds, Fig. 1C. To make tumoroids, following seeding, cells are rolled down to the microwells through the inclined microchannels connected to the microwells. The extra cells in the loading zone are washed by adding culture media to the media reservoir and aspirating through the top microchannels arrays. As shown in Fig. 1D, tumoroids after formation can be embedded inside the relevant hydrogel matrix, mimicking the ECM of the tumor microenvironment. ECM loading is performed through the secondary reservoir of the insert, while the ECM hydrogel solution flows through microchannels, connecting the secondary reservoir to microwells. The ECM hydrogel could be loaded either alone or in mixed with the stromal cell component of the TME, including cancer-associated fibroblast, immune cells, etc. To make the complex tumor model, tumoroids were first generated in the microwell array insert by seeding the cancer cells suspension to the central zone of each quadrant of the insert. Fig. 1E shows the homogenous size of the tumoroids arrays formed within the 4 quadrants of the hydrogel insert. In an additional experiment, the effect of seeding density on the size of tumoroids formed in the insert was characterized. Each quadrant of the insert was dedicated to one seeding density and as depicted in Fig. 1F&G by increasing the seeding number from 50 ×10^3^ to 300 ×10^3^ the tumoroid diameter increased significantly which shows the capability of the insert in controlling the size of the desired formed tumoroids.

**Fig. 1.**
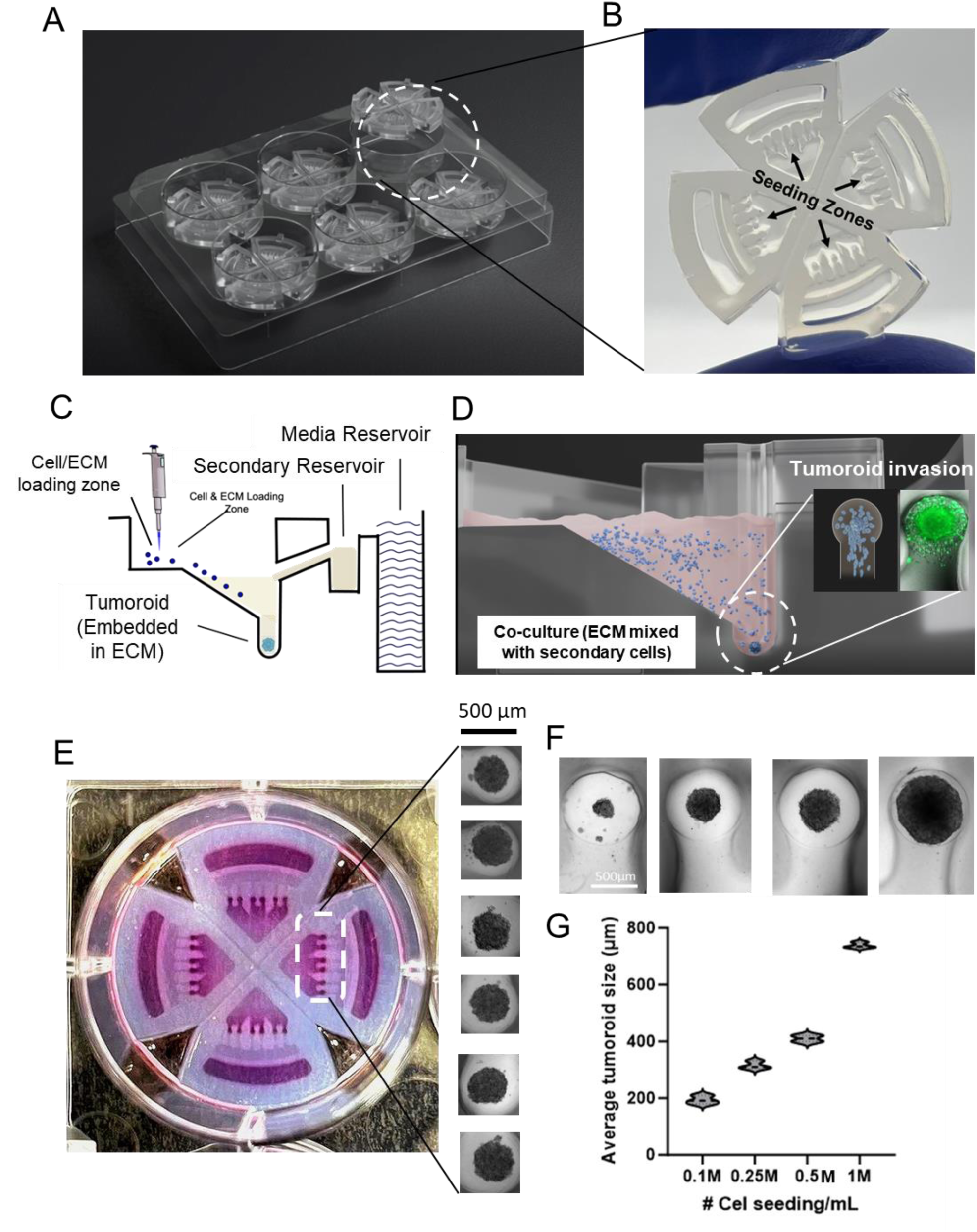
Microfluidic-integrated culture plate for 3D tumoroid array generation. A) Schematic view of the hydrogel inserts in 6-well plate. B) Actual image of the transparent hydrogel inserts with 4 quadrants and an array of addressable microchannel-integrated microwells. C) Side-view Schematic of the tumoroid generation insert. Each insert includes the tumoroid formation and drug delivery modules. D) Schematic view of the tumoroids inside the microwells and guided microchannels after embedding within the ECM for cell migration assay (Inset image). Secondary cells can be embedded inside the cross-linked ECM. E) Top-view of the microwell arrays along with homogenous size of the tumoroids within a quadrant. F) Effect of cell seeding number on the size of the formed tumoroid in the microwells.

We evaluated the ability of our ToP model to keep the glioblastoma cells viable and functional by analyzing the viability and phenotype of the cells cultured in the model. The viability and phenotypic analyses were performed after 72 h of seeding under different cell seeding densities. Fig. S1A illustrates fluorescent images of the U251 tumoroids stained with the Live/Dead kit after 72 h of culture, indicating that the majority of the cells were viable (>95%). The functionality of the cells in the tumor model was also evaluated by the expression of the glial fibrillary acidic protein (GFAP) receptor, a specific marker for astrocytes and glioma cells [41,42]. The U251 tumoroids in the model were immunostained with GFAP after 72 h of incubation and imaged by fluorescence microscopy (Fig. S1B&C). Glioblastoma tumoroids showed a positive GFAP expression, demonstrating the astrocyte phenotype of the cells used in our ToP model.

### 3-2- Screening different GBM-related ECMs and glioblastoma ToP model function

Infiltration of tumor cells into the healthy brain parenchyma is an important feature of GBM malignancy, which makes it almost impossible to resect the tumor through surgery completely. The interaction of membrane proteins (i.e., integrins and cadherin) with extracellular matrix components appears important in mediating the movement and infiltration of tumor cells in healthy brain regions [4]. The Glioblastoma matrix is a complex microenvironment comprising various proteins and polysaccharides. Reelin is an extracellular matrix protein that regulates neural migration during brain development and synaptic plasticity in the adult brain [5]. Reelin canonical signaling is mediated by binding to ApoE2 and VLDLR (Very Low-Density Lipoprotein Receptor) proteins. Reelin can also bind to specific integrins [6]. Within this context, it has been shown that Reelin signaling may modulate GBM growth and substrate-dependent migration [7]. Reelin signaling is decreased in GBM, and Reelin mRNA expression is inversely correlated with malignancy grade [7]. While Reelin expression is decreased in most cancer subtypes studied, it has been shown that high Reelin levels can also be prevalent in other cancer subtypes [8-12]. Our advanced ToP model provides a unique feature that allows for analysis and quantification of tumoroid invasion within the ECM hydrogel without the need to remove or manipulate them after formation in the insert. To investigate the effect of Reelin on the growth and invasiveness of GBM tumor models, we generated arrays of U251 tumoroids in-a-well and they were screened against the varying concentrations of recombinant Reelin protein in culture media. Based on the findings presented in Fig. S2, the investigation into the impact of Reelin on the viability of GBM cells cultured in a 2D environment revealed a slight toxic effect when the concentration of Reelin was increased in the treatment media. Additionally, when the culture time was extended to 48 hours, the toxicity of the 100 nM Reelin was significantly greater than the control condition. Fig. 2A&B demonstrate that by exposing the tumoroids to the same concentration of Reelin from 1 nM to 100 nM, the same toxic trend was observed and the toxicity at 100 nM was reported to be around 27%. Fig. 2A shows the live-dead fluorescence microscopy of the tumoroids treated with different dosage of Reelin in media.

**Fig. 2.**
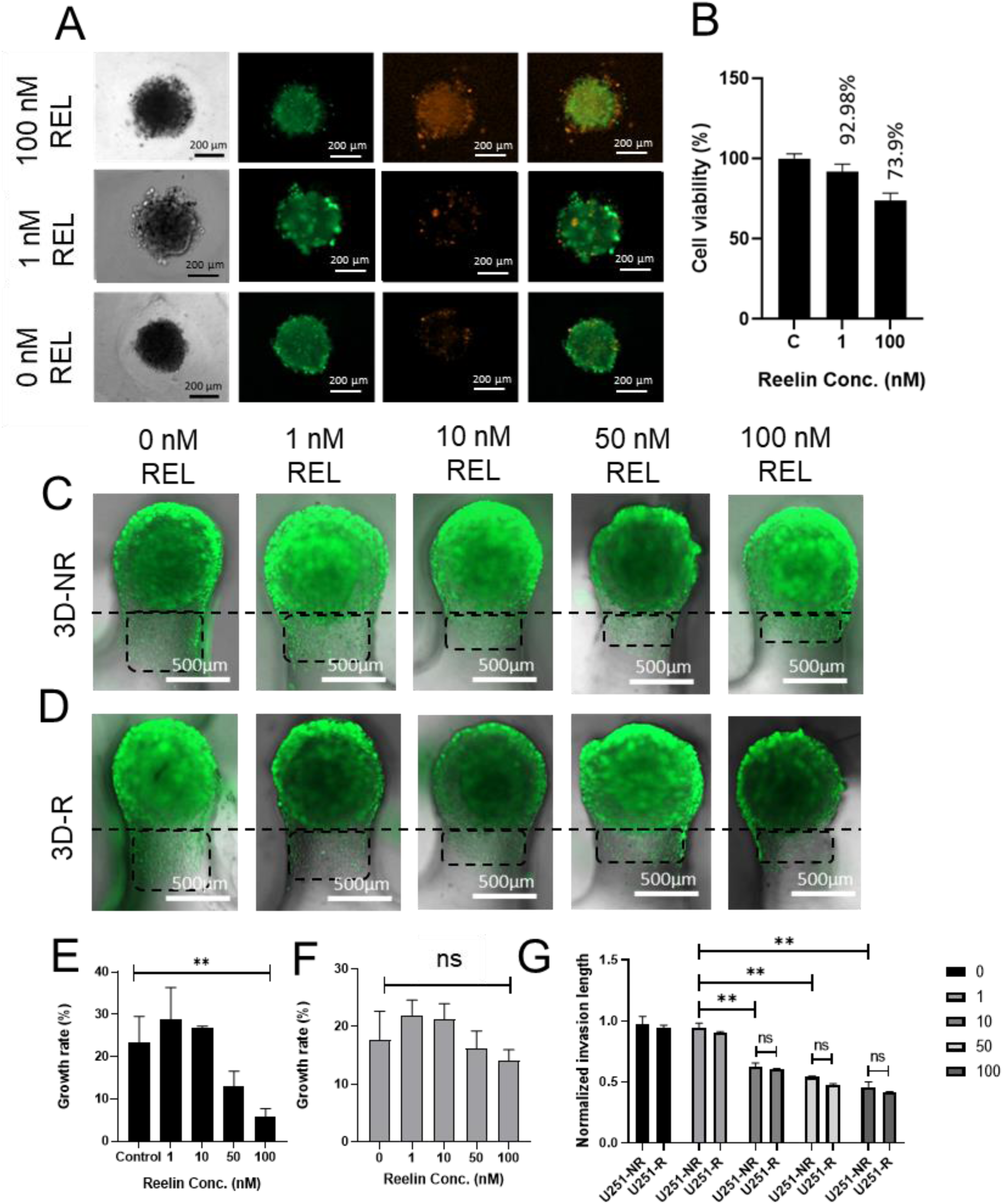
Recombinant Reelin protein affects the growth and migration behaviour of the tumoroids within the ECM in ToP model. A) Live-dead fluorescent images of the tumoroids in the microwells in response to the varying Reelin concentration in culture media. B) Effect of Reelin protein on viability of the U251 tumoroids. Fluorescent microscopic images of the tumoroids within ECM showing the effect of Reelin protein mixed with collagen on invasion behaviour of ECM-embedded C) non-resistant and D) resistant tumoroids. Growth rate of tumoroids exposed to varying Reelin concentrations, E) 3D-NR tumoroids and F) 3D-R tumoroids. G) Quantified normalized invasion length of the tumoroids (3D-R and 3D-NR) in response to the varying Reelin protein mixed with collagen ECM.

In order to mimic the invasion process and infiltration of the tumor cells into healthy brain tissue, we made a collagen solution with a concentration of 4 mg/mL pipetted to the loading zone of the insert while it surrounded the tumoroids in the microwells, Fig1C&D. The tumoroids were then held in place by the buoyancy force of the ECM hydrogel. This study examined tumor invasion in compartmentalized microchannels of the insert. Tumoroids initiated invasion towards the microchannels connected to the microwells. Our results showed that the density of cells and the invasion length along the invasion microchannel compartment increased continuously over time, Fig. S3. Either the cell number per certain area of the channel or the normalized invasion length can be used for the quantification of invasion. Here, we quantified the invasion by counting the length of the of cells migrated within the ECM. Fig. S3 shows the progression of the number of invaded cells every 24 h. The invasion length was normalized with the diameter of the tumoroid before invasion and immediately after ECM embedding.

In order to study how Reelin conditioned ECM affects the growth and invasion of tumoroid models, we mixed Reelin with the primary collagen solution in equal concentrations. This mixture was then added to the tumoroids in the well. After surrounding the tumoroids for an hour at 37 C, the Reelin-loaded collagen hydrogel was cross-linked, and we measured the diameter of the tumoroid over time. As depicted in Fig. 2C&D. tumoroids were able to invade with different lengths through microchannels connected to microwells within the insert design, after exposure to different ECM conditions. Fig. 2E shows that reelin-conditioned ECM has a significant effect on the growth rate of the tumoroids, so by increasing the Reelin concertation in collagen from 1 to 100 nM, the growth rate is decreased to less than 10%. Growth rate in this analysis was calculated based on the primary tumoroid size of each condition right away after the ECM embedding and cross-linking.

Parallel to the tumoroid growth measurement in the Reelin conditioned ECM, we observed a descending trend for the normalized invasion length of the tumoroids within the hydrogel with increasing the amount of Reelin in the collagen ECM. Increased Reelin concentrations in the collagen hydrogel from 1nM-100 nM make glioma tumor cells more invasive and more likely to migrate within the ECM. The higher invasive behavior of the glioblastoma tumoroids in the absence of the recombinant Reelin in ECM hydrogel is attributed to the silencing of the Reelin and its downstream effector Dab1 gene expression in malignant tumors which is inversely correlated with the malignancy and invasiveness of the tumors ^25,26^.

To investigate the impact of Reelin on tumor invasiveness, we conducted experiments using a alternative version of the GBM tumoroids in our ToP model. We utilized U251 glioma tumoroids that have intrinsic resistance to chemotherapy. These tumoroids, called 3D-RT, were generated by exposing the cancer cells to 250 uM concentration of TMZ. The chemoresistance U251 cell lines were established and characterized previously in our group^27^. To investigate the chemosensitivity of the established cell lines in 3D microenvironment, non-resistant and TMZ-resistant tumoroids were formed in our microfluidic-integrated culture plate. As depicted in Fig. S4, TMZ-resistant tumoroids (3D-RT) displayed significantly lower sensitivity to TMZ (250 µM) as compared with non-resistant spheroids (3D-NR), Fig. S4. In order to study how TMZ-resistant tumoroids grow and spread in Reelin-conditioned ECM, we embedded 3D-R tumoroids in a collagen ECM with different concentrations of Reelin, similar to the experimental procedure with 3D-NR tumoroids. As depicted in Fig. 2 D&G, we observed the same invasive behavior in 3D-R tumoroids as 3D-NR ones, where the normalized invasion length of the tumoroids significantly dropped at 10 nM Reelin-conditioned ECM, and it was followed by non-significant reduction by increasing the Reelin dosage to 100 nM. Moreover, as shown in Fig. 2F, the growth rate of the 3D-R tumoroids followed a similar pattern where the growth rate decreased with an increase in Reelin concentration up to 100 nM. However, there was no significant difference between the control and the 100 nM conditions. Interestingly, the growth rate of the 3D-RT tumoroids at 100 nM Reelin concentration was reported to be above 10%, which is higher than that of 3D-NRT tumoroids. This reflects the less sensitivity of the chemoresistance U251 tumor cells to the inhibitory effect of Reelin in the GBM tumour microenvironment. Based on our data, it appears that the influence of reelin on GBM invasion and progression behavior could be a significant factor in regulating GBM behavior. This finding can spark further interest in exploring the potential of the reelin pathway as a target for GBM treatment.

### 3-3- Chemotherapeutic screening of the GBM tumoroids in single and multi-drug treatment settings

To investigate the clinical response of the Reelin conditioned ECM within the 3D glioma in vitro model to the chemotherapeutic treatment of the GBM, we screened the varying dosages of TMZ as a clinically FDA approved chemo pro-drug for the GBM patients against an array of tumoroids on-a-plate. Tumoroid response to TMZ treatment in the presence and absence of the Reelin in ECM was studied through invasion inhibition, viability measurement of the cells within the tumoroid and tumoroid size reduction analysis, Fig. 3. In order to assess how Reelin and TMZ affect the invasiveness of tumoroids, we measured the relative invasion length of the tumoroids when exposed to different concentrations of TMZ. Fig. 3A&B shows that under both ECM conditions, there is a gradual reduction in invasion length of the tumoroids as the concentration of TMZ increases. However, the decrease in relative invasion length is more pronounced in the Reelin ECM compared to collagen. Additionally, the number of cells that invaded through the microchannel in each treatment condition was measured by counting them within a certain area of the microchannel, Fig. 3C. As shown in Figure 3D, the viability of tumoroids is more negatively affected by TMZ when embedded in Reelin-conditioned ECM compared to collagen ECM. This difference in viability is particularly evident at higher dosages of TMZ, where a significantly higher level of toxicity is observed in tumoroids embedded in Reelin ECM compared to those embedded in normal ECM at a concentration of 500 µM. These results confirm the slight toxic effect of Reelin on the viability tumoroids that was demonstrated in Fig. 2A. To determine the inhibitory effect of TMZ in the presence of Reelin-conditioned ECM on tumoroid progression, we measured tumoroid size reduction on day 3 of the treatments. In Fig. 3E-G, a significant reduction of approximately 20% and 30% in tumoroid size was observed after exposure to 500 µM TMZ in collagen and collagen-Reelin ECMs, respectively, compared to the lower dosages. However, at 500 µM TMZ concentration, the Ccollagen-Reelin ECM had a greater size reduction effect than the collagen ECM, indicating the inhibitory effect of Reelin in ECM was abolished.

**Fig. 3.**
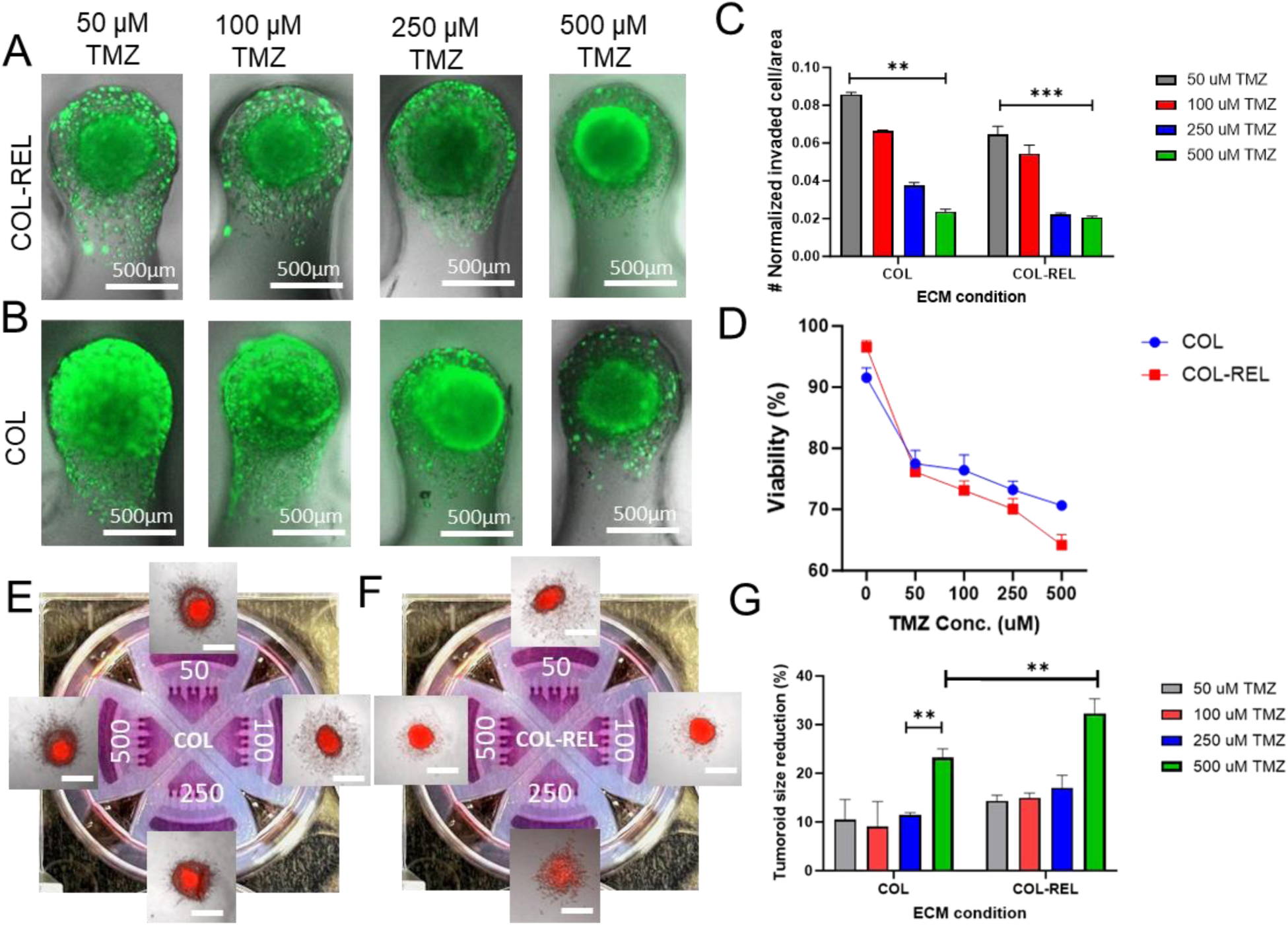
GBM ToP model is used for screening the effect of ECM composition in different TMZ dosages on U251 tumoroids. Fluorescent microscopic images of the tumoroids embedded in A) Collagen-Reelin and B) Collagen in response to the increasing concentrations of the TMZ. C) Quantified normalized invasion length of the tumoroids in response to the different ECM conditions and varying TMZ treatment dosages. D) Cell viability curve of the tumoroids embedded in 2 different matrixes exposed to the varying TMZ dosages. E, F) microscopic images of the treated tumoroids in each ECM and drug condition, along with the G). Size reduction percentage of the tumoroids exposed to the varying TMZ dosages in collagen and collagen-Reelin ECMs. The scale bar is 500µm.

The current results showed that the inhibitory effect of Reelin on invasion behavior of the tumoroids within the ECM is more dominant in lower concentration of TMZ while this effect is less significant on 500 µM TMZ where there is no difference between invasion density of the tumor cells in collagen and Reelin-collagen ECMs. The effect of Reelin on invasion inhibition of tumor cells has been studied for brain tumor and breast cancer and it is shown that Reelin gene, which encodes the extracellular glycoprotein Reelin is downregulated in these cancer types^25,28^. Despite TMZ is usually suggested as the first line of chemotherapy treatment on malignant glioma cancers, our physiological in-vitro model demonstrated that exposing the tumoroids to a high dosage of TMZ (500 µM) didn’t show any significant toxic effect on their viability and, therefore to achieve efficient toxicity on the tumoroids, higher concentrations of TMZ are necessary. However, increasing the dosage of TMZ will have negative side effects on the viability of healthy glioma and astrocyte cells in the brain tissue^29^. Recently, combination chemotherapy has been suggested as a new approach to improve the efficacy of treating solid tumors^7^. This is achieved by exposing the tumors to a combination of drugs, resulting in a potential synergistic effect. In this regard, we studied the effect of multidrug treatment on GBM tumoroid progression and invasion. We exposed arrays of U251 tumoroids embedded in 10 nM Reelin conditioned ECM to varying concentrations of TMZ, ranging from 50 µM to 500 µM, following treatment with an iron chelators drug, deferoxamine (DFO). Iron can play a dual role by participating in both promotive/inhibitive reactions related to ROS generation, DNA synthesis, etc. Hence, Iron chelators can remove iron from the cell and deprive cancer cells of their highly required source, and hence decreasing viability and proliferation ^30^. In this regard, Iron chelating drugs such as DFO have attracted attention as potential therapeutic drugs through tumor-suppressing effects by causing iron depletion^31^. A study on the effects of DFO on tumor viability and progression in the collagen ECM was conducted using the same platform. Results showed that DFO can cause a toxic effect of about 45% on 3D tumoroids in a well^27^. The current study investigated the combination of DFO and TMZ treatment in a sequential manner within the Reelin-conditioned ECM.

The effectiveness of a combined treatment approach for GBM tumors was confirmed through cell death analysis and measurement the viability of the cells. This was further supported by a reduction in tumoroid size, as observed in the microscopic analysis. According to Fig. 4A-C, when tumoroids were treated with 100 µM DFO for three days followed by 100 µM TMZ treatment, their viability dropped to less than 50% compared to the same treatment of 100 µM TMZ combined with 10 µM DFO. Moreover, metabolic activity assay results of treated tumoroids in Reelin conditioned ECM, showed that combination of 100 µM DFO and 100 µM TMZ in sequential exposure, caused more toxic effect in comparison to the highest concentration of TMZ (500 µM) in single treatment experiment. The study found that the increasing TMZ concentration with the presence of DFO led to a significant reduction in the viability of tumoroids. The IC50 value was achieved at a condition where TMZ concentration was 100 µM and DFO concentration was 100 µM. However, when the tumoroids were treated with lower primary iron chelator doses, the combinatorial effect was less significant in achieving the IC50 value in binary drug therapy so that the IC50 value of TMZ increased to 250 µM. According to Fig. 4D, the size reduction results of the tumoroids after binary drug treatment are in line with the toxicity data, and the binary treatment with a higher concentration of DFO (100 µM) resulted in significantly greater size reduction than the same TMZ treatment combined with 10 µM DFO. Interestingly, the binary treatment at a lower iron chelator dose (10 µM) didn’t show any significant change in tumor size reduction rate compared to the single treatment with the same TMZ range. However, after the concentration of DFO was increased to 100 µM, the rate of reduction in tumoroid size was found to be more significant in the binary treatment as compared to the single TMZ treatment. This resulted in an increase in the size reduction percentage of 40%, 53%, 51%, and 21% in binary treatment for the TMZ treatment range from 50 µM to 500 µM, when compared to the same single TMZ therapy range. The invasion behavior of tumoroids in response to varying concentrations of the combination therapy was also quantified in Fig. 4E&F. While relative invasion length of the tumoroids with IC50 of 10 and 100 µM DFO did not show significant differences with lower TMZ concentrations, the invasion analysis revealed that the 1000 µM TMZ and 100 µM DFO condition had the greatest effect in decreasing the invasion length of the tumoroids within the ECM.

**Fig. 4.**
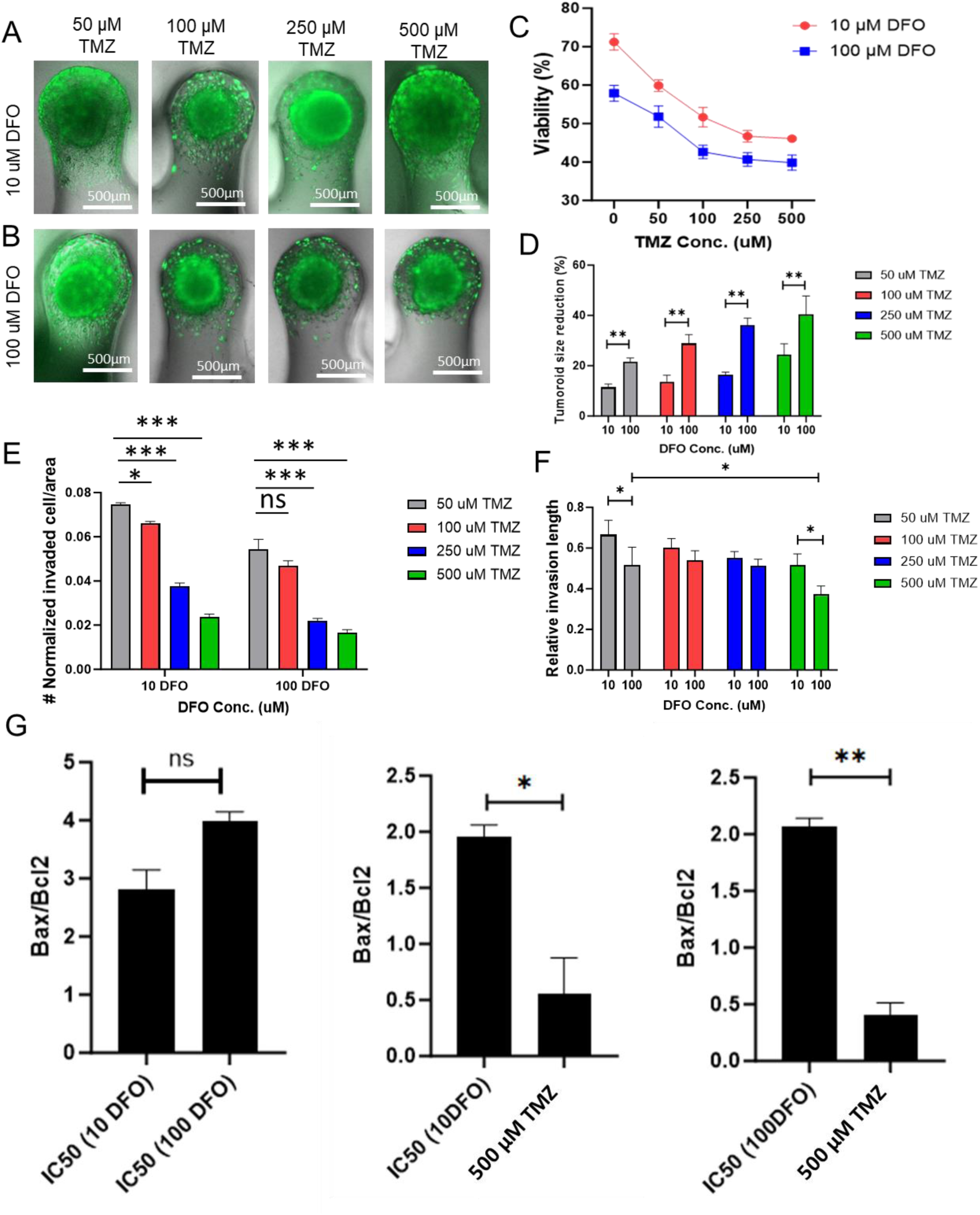
Combinatorial drug screening on the ToP GBM models. Fluorescent microscopic images of the tumoroids embedded in Collagen-Reelin matrix in response to varying concentrations of TMZ following treatment with A) 10 µM DFO and B) 100 µm DFO. C) Viability trend of the tumoroids embedded in Collagen-Reelin ECM exposed to the varying TMZ dosages after DFO treatment. D) Size reduction percentage of the tumoroids in collagen-Reelin ECM exposed to the varying TMZ and DFO dosages. E) Quantified normalized invaded cells/microchannel are of the tumoroids and F) relative invasion length in response to the different ECM conditions and varying TMZ treatment dosages. G) Bax/Bcl2 expression of the tumoroids at IC50 dosage of DFO treatments with 10 and 100 µM concentrations.

To explore the cell death mechanism of the tumoroids after exposure to the combination of DFO/TMZ compared to the TMZ single treatment, expression of Bax and Bcl-2 in IC50 of the TMZ in binary therapy and single TMZ therapy (500 µM) were investigated using real-time PCR analysis. The ratio of Bax to Bcl-2 acts as a switch for cell death, determining if cells will survive or die in response to an apoptotic stimulus. When the Bax/Bcl-2 ratio increases, the cell’s resistance to apoptosis decreases, resulting in increased cell death and higher toxicity. As shown in Fig. 4G, the ratio of Bax/Bcl-2 expression in IC50 of the 100 µM DFO treatment was reported to be more than IC50 of 10 µM DFO treatment in a non-significant way which indicated the prominent role of iron chelator in induction of apoptosis in the treated tumoroids. On the other hand, by comparison of the apoptosis induction ability of the single TMZ treatment and its combination with 10 µM and 100 µM DFO treatment, we found that the presence of the DFO in the treatment regime of the tumoroids has a significant effect on apoptosis stimuli to induce cell death of the GBM tumoroids.

### 3-4- Tumor stroma incorporation in pancreatic ToP in vitro model

The tumor microenvironment, which includes cancer-associated fibroblasts, endothelial cells, and tumor-associated immune cells, plays a significant role in the initiation, progression, and migration of tumor cells^32^. In the previous section, we showed that our advanced tumoroid-on-a-plate in vitro model can incorporate customized ECM for relevant cancer progression assays. It also enables the creation of a complex TME, which allows for the study of stromal tissue impacts on cancer progression and invasive behavior after tumoroid formation on the same plate. In the current study, we showed the capability of co-culturing under two different methods of direct and in-direct 3D co-culture. Pancreatic tumors often exhibit a dense stromal microenvironment characterized by abundant stromal cells, including pancreatic tissue fibroblasts and pancreatic stellate cells that contribute to producing and depositing fibronectin and collagen I and III ^33^. These stromal cells play a critical role in the progression and aggressiveness of pancreatic tumors ^34^. Additionally, they create a physical barrier around cancer cells, making them less accessible to chemotherapy and other therapeutic interventions^35^. Furthermore, stromal cells can induce epithelial-mesenchymal transition (EMT) in cancer cells, promoting their invasive behavior and resistance to apoptosis. Understanding the complex interactions between stromal cells and pancreatic tumors is essential for developing targeted therapies aimed at disrupting the tumor-stroma crosstalk and improving treatment outcomes for pancreatic cancer patients. In this regard, we used our microfluidic-integrated culture plate insert to make a co-cultured pancreatic tumor model and investigate the role of human-derived fibroblast (HDF) cells on formation, invasive behavior and drug response of the Panc-1 pancreatic tumoroids in collagen ECM.

As shown in Fig. 5A, co-cultured pancreatic tumor model was accomplished by co-seeding the Panc-1 cells mixed with certain concentration of the fibroblast cells in culture media to make the tumoroids with different cancer/fibroblast ratios. Panc-1 cell suspension with the density of 200,000 cells in 50 µL of the media, tumoroids were formed after 3 days. However, growth of the 3D co-culture tumoroids was monitored over time till day 7 using bright filed microscopy Fig. 5B.

**Fig. 5.**
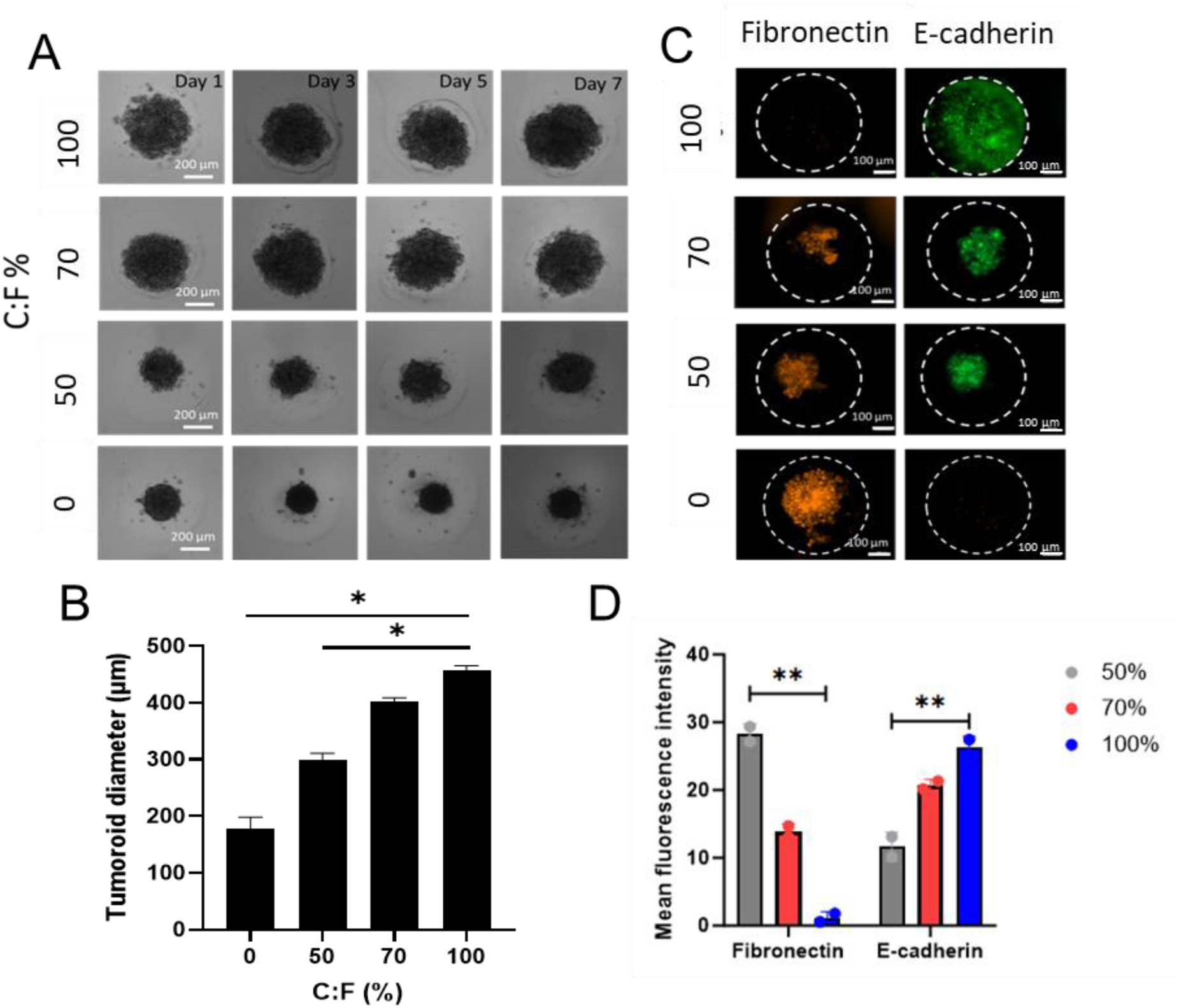
Co-cultured Pancreatic adenocarcinoma ToP model recapitulates the stromal effect of TME. A) Bright-field images of the Panc-1-HDF co-cultured ToP models with different ratios of cancer to fibroblast cells. B) Quantification of the co-cultured tumoroid diameter with different cancer-to-fibroblast ratios. C) Immunostained fluorescent images of the E-cadherin and fibronectin in co-cultured tumoroid diameter with different cancer-to-fibroblast ratios. D) Quantification of mean fluorescent intensity of the E-cadherin and Fibronectin markers in each co-culture condition.

To investigate the effect of direct co-culture (co-seeding) on the behavior of the tumoroids, it is depicted in Fig. 5B that by decreasing the ratio of the cancer cells to fibroblast cells in the primary cell suspension, the tumoroids are shrunk. This size change is attributed to the effect of fibroblast contribution in co-cultured tumoroid formation so that fibroblast cells induce a contractile tension to the multi-cell aggregates within the body of the tumoroids and make that size smaller and maximizing the contact between cells ^36,37^. In Fig. 5B, we observed a reduction of over 50% in the size of cell aggregates in microwells, from 100% cancer cells to 0% cancer cells. It is important to note that the chosen ratio of cancer cells to fibroblasts (C/F) in the current experiment demonstrates the ability of the current platform to create co-cultured models with varying ratios of stromal components, depending on the type of cancer and the ultimate goals of the model. These data are confirmed by the in-well florescent staining of the epithelial tumor cells and fibroblast cells with their specific markers E-cadherin and fibronectin respectively. As illustrated in Fig. 5C&D, the accumulation of the cancer cells expressing E-cadherin is close to the center of the tumoroids by increasing the population of the fibroblast cells to 50%. E-cadherin is a cell adhesion molecule necessary for the maintenance of intercellular contacts and cellular polarity in epithelial tissues, and is a key feature of EMT, showing the invasive behavior of the cancer cells ^38,39^. As measured in Fig. 5D, by increasing the number of the fibroblast cells in the co-cultured tumoroids, a decrease in E-cadherin expression of the immunostained panc-1 tumor was reported, characterized by fibroblast activation.

In the next step, we used the co-cultured pancreatic ToP model to screen the tumoroid response against chemotherapeutic agents in single and binary settings. The current 3D culture plate insert allows for four experimental conditions per insert. In order to screen the response of co-seeded pancreatic tumoroids to a range of chemo drugs, we treated them with a combination of gemcitabine (GEM) and DFO in a sequential manner. This was done in a similar way to the glioblastoma tumoroids in the previous section. During the last two decades, gemcitabine (GEM), a nucleoside analog of deoxycytidine, has been the standard chemotherapeutic agent for pancreatic cancer ^40,41^. Recently, new combination chemotherapies, such as regimens combining fluorouracil, irinotecan, oxaliplatin, and leucovorin (FOLFIRINOX) ^42,43^ have been reported for treating PDAC. However, while combination chemotherapies have shown therapeutic advantages over single-agent GEM, they also have a high incidence of side effects. To design an effective binary therapy against the PDAC with lower off target side effects, we studied the combination of GEM and the iron chelator drug DFO with a varying concentration to screen their toxic effect on the viability co-cultured pancreatic tumoroid model. According to literature iron chelator compounds can suppress movement and inhibit invasion of pancreatic cancer cells in addition to their antiproliferative activity in a dose-dependent manner^44^. Figure 6A shows the combinatorial drug treatment design on our ToP model. It also displays the normalized fluorescent intensity of the stained live and dead cells’ population within the tumoroids under each treatment condition. As shown in Fig. 6 A&B, by increasing the concentration of GEM in single treatment regime, the fluorescent intensity of the live cells population significantly drops at 100 µM GEM and stays consistent at 1000 µM GEM. On the other hand, when we combined DFO with GEM, we observed a synergistic effect at 10 µM DFO along with 100 µM GEM. In this combination, the intensity of live cells within the tumoroids was lower compared to the single GEM treatment, and the population of dead cells showed higher red intensity compared to other treatment conditions. However, the fluorescent live-dead image array depicted in Fig. 6B only focuses on dead cells in the tumoroid body, but it does not consider detached dead cells followed by the washing and staining steps. To get a more accurate understanding of the viability of the tumoroids in each treatment combination, the metabolic activity analysis was conducted using presto blue assay (determining NADH levels) on tumoroids after exposing to each treatment condition for 3 days. As shown in Figure 6C, we observed a similar quantified trend in viability of the co-cultured tumoroids under different treatment conditions while the IC50 of the binary treatment was reported at 10 µM DFO, 100 µM GEM condition.

**Fig. 6.**
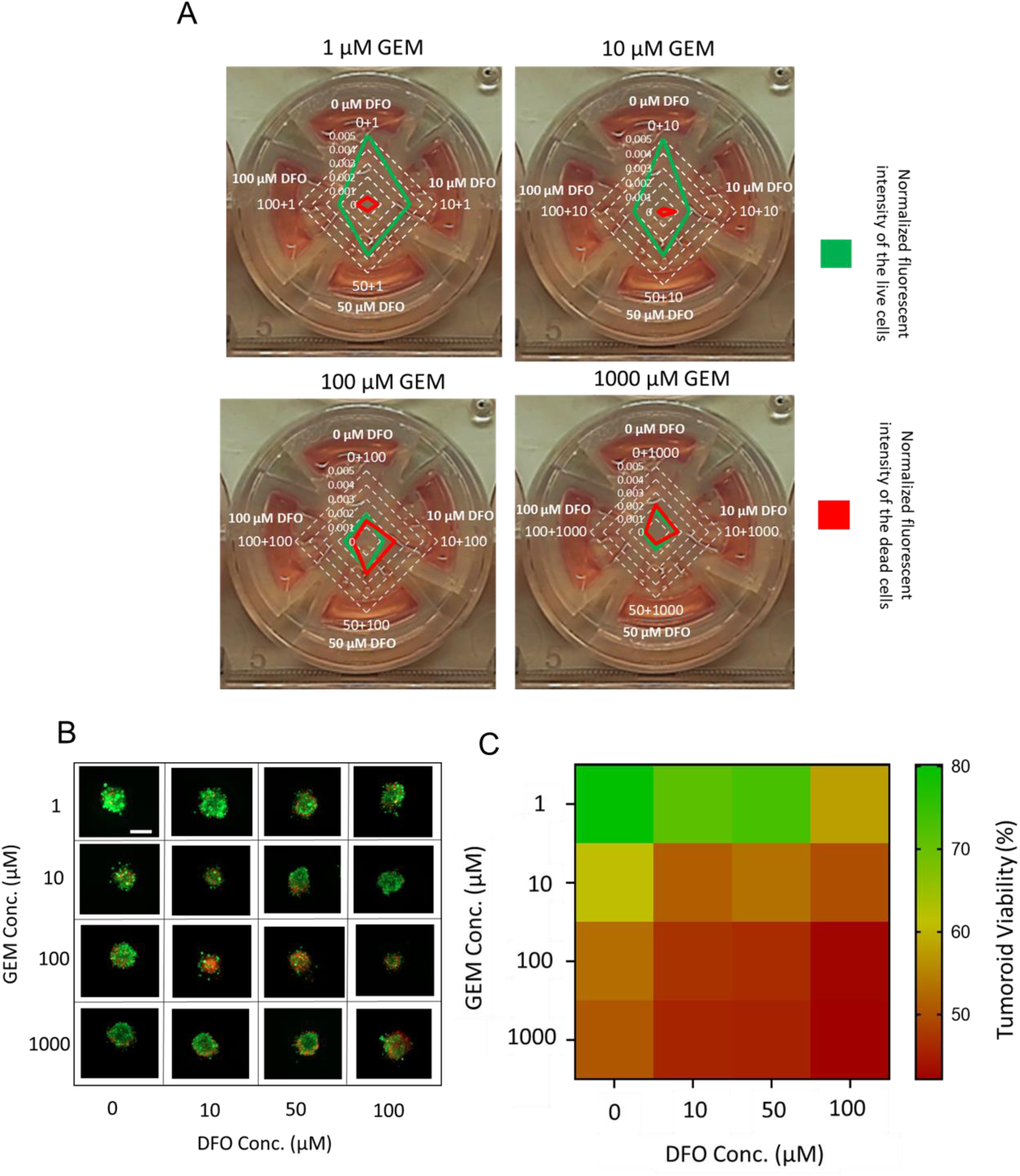
Combinatorial drug screening on the ToP pancreatic co-cultured models. A) Normalized fluorescent intensity of the live and dead cells population within the treated tumoroids with different combinations of DFO and GEM dosages. B) A fluorescent image array showing the viability of ToP models screened under various treatments. C) Heat-map analysis of the tumoroid viability under different treatment combinations.

In order to assess the impact of direct co-culture on the invasion behavior of pancreatic tumoroids within the ECM, we selected C/F ratios of 100% and 50% for co-cultured tumoroids in our model and analyzed the length of invasion in each condition. Our findings depicted in Fig. 7A-C, demonstrate that fibroblast cells have a significant effect on increasing the invasion length of tumoroids in our model. Specifically, we observed that increasing the population of fibroblasts to 50% resulted in a normalized invasion length exceeding 100% compared to panc-1 tumoroids on both day 1 and day 5 of the experiment. Moreover, in Fig. 7 B&D, we showed that by increasing the ratio of the fibroblast to cancer cells in our ToP model followed by ECM embedding, the same trend of E-cadherin expression was reported for the co-cultured tumoroids so that, at 50% fibroblast we observed more invasive behavior and less E-cadherin expression in the tumor cells within the model. Generally, it was observed that fibroblast cells affected tumor cells in the co-culture model leading to the induction of the Epithelial-mesenchymal transition process in tumor cells ^45^.

**Fig. 7.**
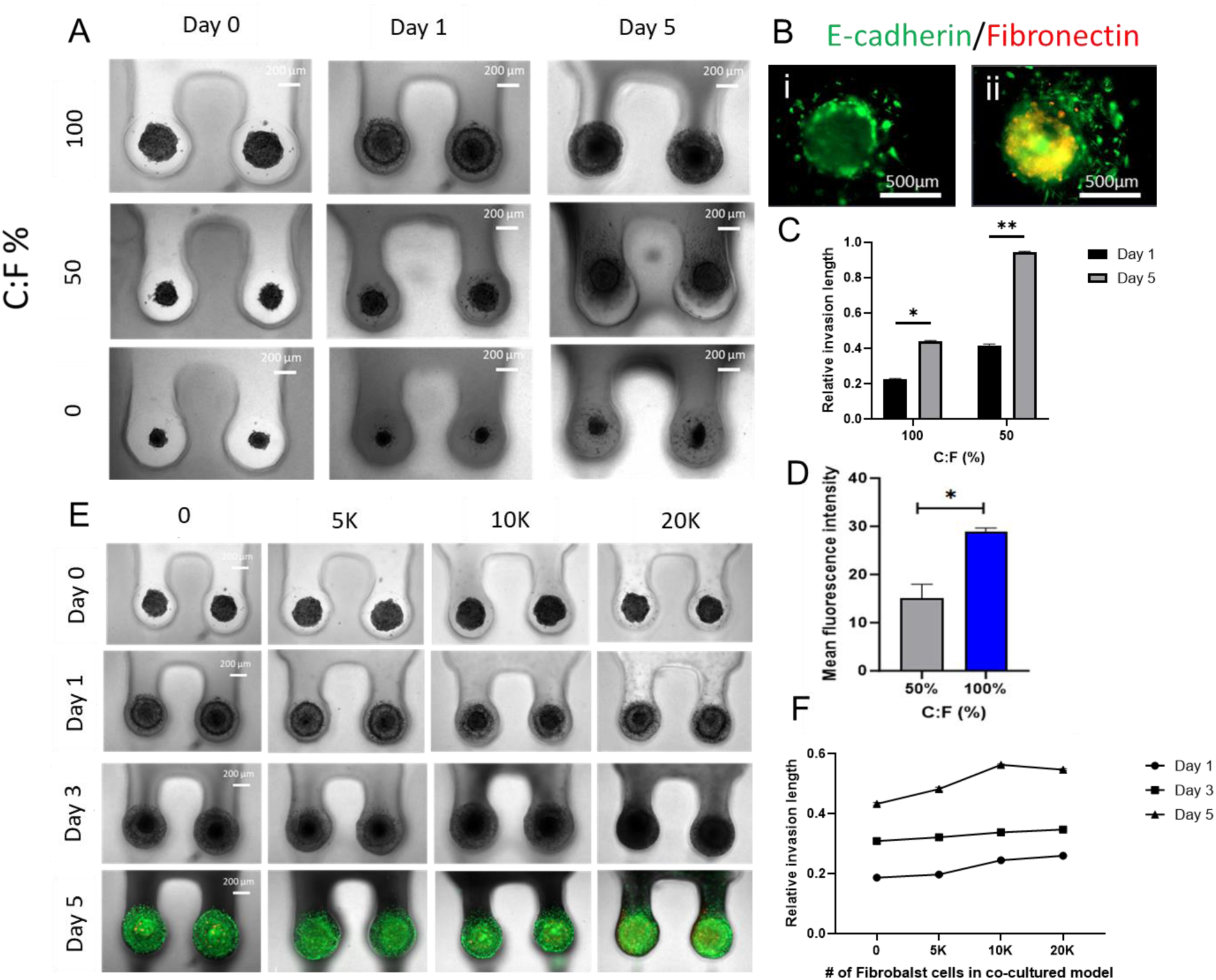
Direct and in-direct co-culture ToP model predicts the invasion behavior of the PDAC tumoroids. A) Bright field microscopic images of the co-seeded (direct co-cultured) Panc-1 and human-derived fibroblast cells invaded within the ECM. B) Immunostained fluorescent images of E-cadherin (green) and Fibronectin (red) expression in co-cultured tumoroids with C:F ratio of i)100% and ii) 50%. C) Relative invasion length analysis of the direct co-cultured tumoroids with C:F ratios of 100% and 50 %. D) Mean fluorescent intensity measurement of the E-cadherin in direct co-cultured tumoroids with C:F ratios of 100% and 50 %. E) Bright field and fluorescent live-dead microscopic images of the in-direct co-cultured Panc-1 and HDf cells invaded within the ECM with different number of the HDF cells/quadrant of the insert. F) Relative invasion length analysis of the indirect co-cultured tumoroids with varying fibroblast numbers.

Regarding the capability of our advanced 3D culture plate in in-situ ECM embedding of the tumoroids in the microwell, there is an opportunity to conduct the co-culture using the in-direct method as well. In this method, we embedded the tumoroids in microwell with mixture of the collagen solution and certain number of fibroblast cells. To capture the in vivo effect of fibroblast cell number in the stromal tissue of the pancreatic tumoroid on the invasive behavior of cancer cells, we mixed the increasing number of fibroblasts form 5000 cells to 20000 cells with collagen ECM hydrogel and loaded to the ECM loading zone of each quadrant of the insert, Fig. 7E. Invasion behavior of the tumoroids in the presence of the fibroblast loaded ECM cells was monitored under brightfield and fluorescent microscopy till day 5, Fig. 7E.

As depicted in Fig.7E&F, the analyzed data regarding the invasion of tumoroids in different conditions confirmed the previous findings of the direct co-culture experiment. It was observed that as the number of fibroblast cells increased in the co-cultured model, the tumoroids exhibited a more invasive behavior, and had a higher invasion length within the matrix, on each day of culture from day 1 to day 5.

The findings in the current study demonstrate the potential of using tumoroids on a plate as an all-in-one 3D culture insert for various applications in human-mimicked tumor organoid modeling, as well as ECM and drug screening. This approach can help in better understanding the mechanisms underlying cancer growth and progression, as well as how tumors respond to pre-clinical therapeutic interventions.

## 4- Conclusion

Developing drugs involves testing different therapeutic compounds on cells grown in mono-layer cultures and animal models. This is followed by multiple stages of clinical trials, which can take several years and cost billions of dollars. Unfortunately, the current tissue models used in drug development have a failure rate of over 95% to bring that drug to therapy, making it challenging to predict how humans will react to the drugs accurately. Additionally, this approach results in a significant loss of animal lives. To address these issues, we aimed to create a fast and cost-effective 3D tumoroid culture platform that is compatible with standard well plates. This platform also benefits from compartmentalized sections with addressable microwells for TME involvement and drug screening assays. The developed ToP mimics the complexity of the TME and provides a human-mimicked platform for evaluating the impact of different treatment approaches on tumor cell progression and invasion. Our study showed that the 3D ToP model can evaluate the impact of Reelin as an abundant brain ECM composition on the invasion and progression of GBM tumoroids. Furthermore, the GBM ToP model can screen the array of tumoroids embraced in the tumor-mimicked ECM against single and multi-therapeutic agents. The results demonstrated the effectiveness of the combination therapy on growth and invasion rate inhibition of the GBM tumoroids on-a-plate. Additionally, we demonstrated that the current ToP model could involve the stromal component of the TME in compartmentalized microchannels without needing to manipulate the tumoroid after its generation on the plate. Our results showed that the co-cultured PDAC tumoroid model with human-derived fibroblast cells expresses greater invasive behavior and less E-cadherin expression, representing the EMT process in the tumor cell phenotypes. Furthermore, the co-cultured ToP model evaluated the effect of new synergistic therapeutics on cell death and progression of solid tumors by screening the combination of the iron-chelator DFO and GEM as a common PDAC chemo drug.

## Acknowledgments

We would like to acknowledge the Natural Sciences and Engineering Research Council of Canada (NSERC), the Canada Foundation for Innovation (CFI), and the BC Knowledge Development Foundation.

## Notes

### Competing Interest Statement

The authors have declared no competing interest.

