## supplementary Information for "Tumoroid-on-a-Plate (ToP): Physiologically Relevant Cancer Model Generation and Therapeutic Screening"

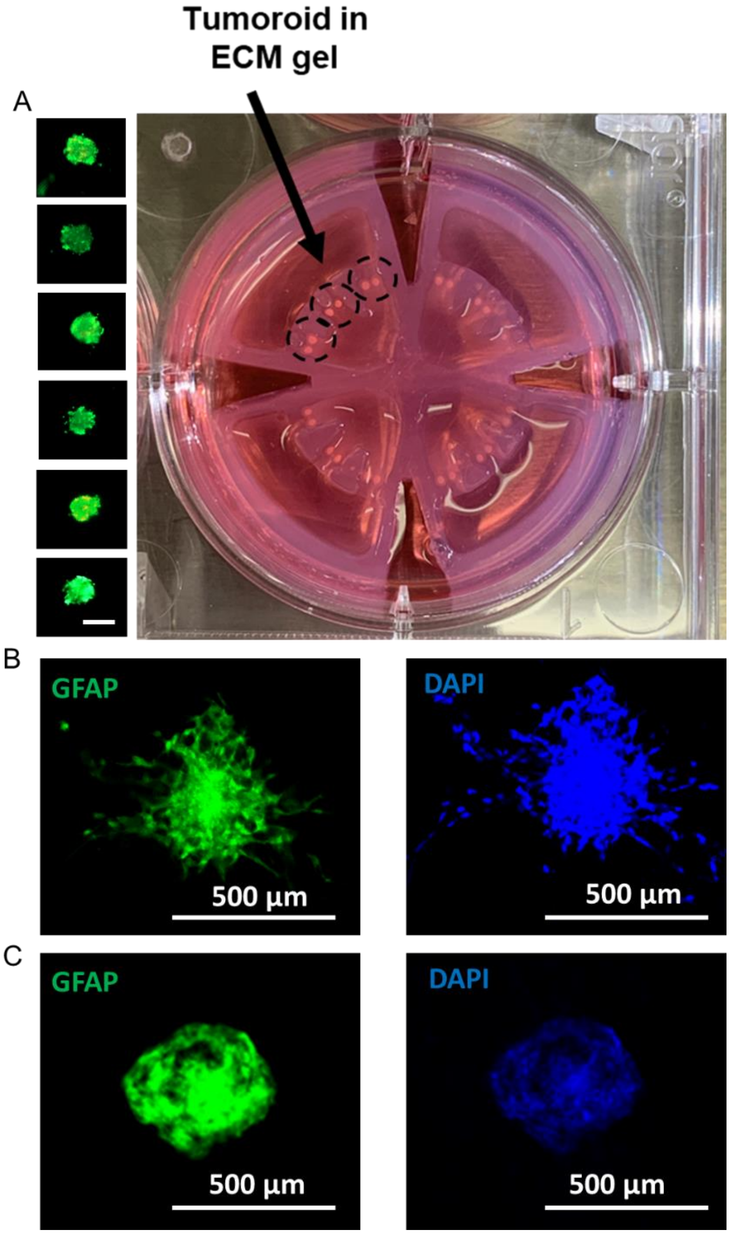


Fig. S1. A) Live-dead fluorescent images of the GBM tumoroids on-a-plate along with the actual image of them in the hydrogel insert. The scale bar is 500 µm. Fluorescent image of GFAP expression in GBM tumoroid B) embedded in Reelin-collagen ECM and C) bare conditions.

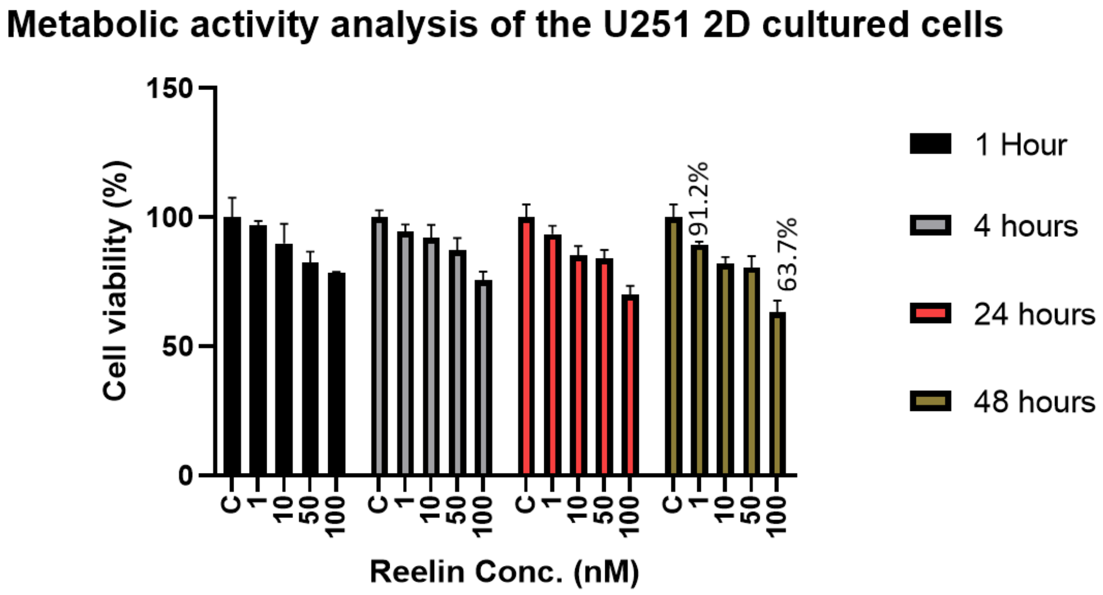


Fig. S2. Viability of 2D cultured U251 GBM cells after exposure to the recombinant Reelin protein in varying concentrations.

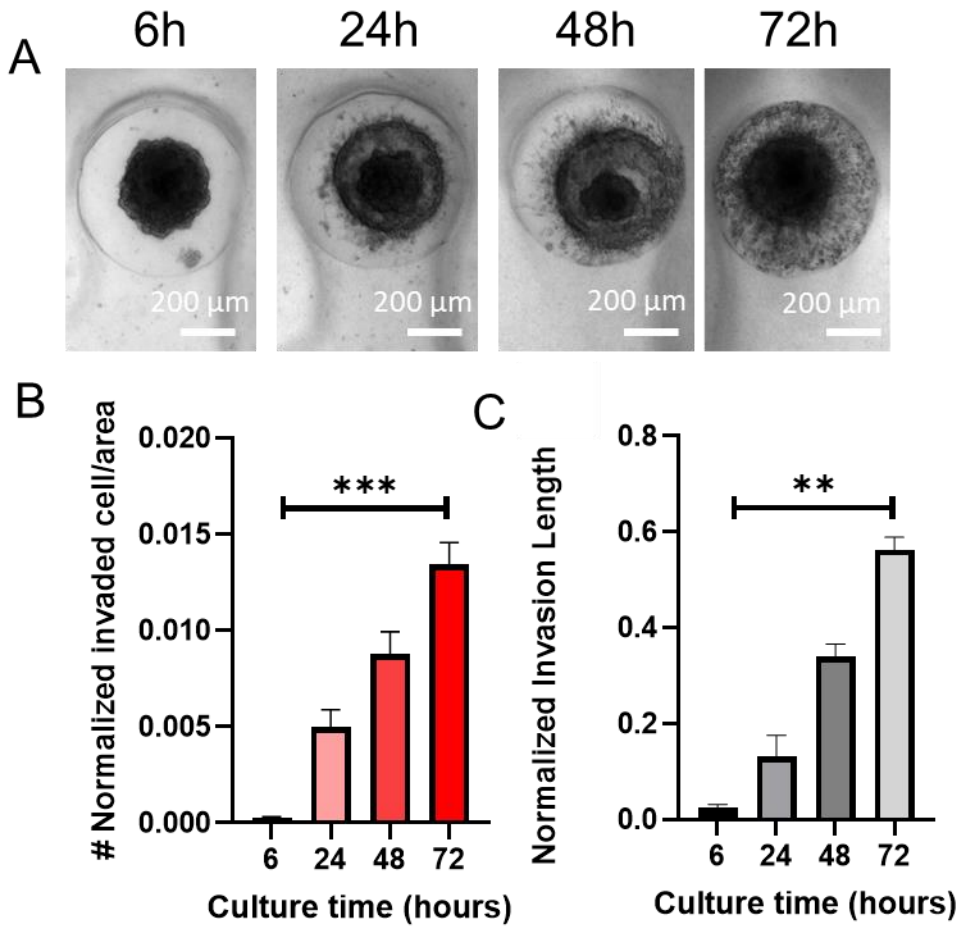


Fig. S3. A) Bright filed images of the GBM tumoroid invasion on-a-plate. B,C) Normalized invaded cell/are and invasion length of the GBM tumoroids within the collagen ECM on-a-plate.

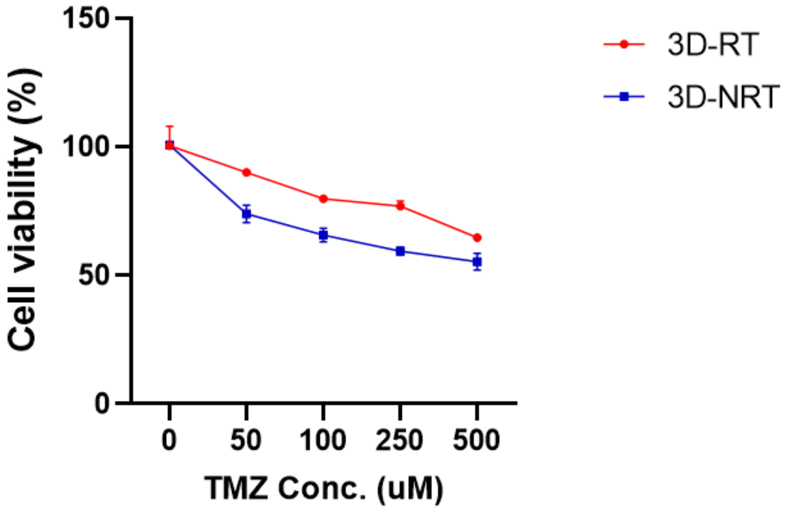


Fig.S4. Cell viability curve of the TMZ-resistant (3D-RT) and non-resistant tumoroids (3D-NRT) exposed to the varying doses of TMZ.
